# Action planning and affective states within the auditory peripersonal space

**DOI:** 10.1101/2020.08.04.236380

**Authors:** Mehrdad Bahadori, Roberto Barumerli, Michele Geronazzo, Paola Cesari

**Affiliations:** Department of Neurosciences, Biomedicine & Movement Sciences, University of Verona, 37131 Verona, Italy; Department of Information Engineering, University of Padova, 35131 Padova, Italy; Department of Architecture, Design, and Media Technology Aalborg University, 2450 Copenhagen, Denmark

**Keywords:** peripersonal space, action planning, affective states, emotion perception, distance estimation, cochlear implants

## Abstract

Fast reaction to approaching stimuli is vital for survival as for sounds entering the individual auditory Peripersonal Space (PPS). Closer sounds have found to provoke higher motor cortex activation particularly for highly arousing sounds, showing the close relationship for perceptual components of the sounds and motor preparation. Here Normal Hearing (NH) individuals and Cochlear Implanted (CI) individuals have been compared in their ability to recognize evaluate and react to affective stimuli entering the PPS. Twenty (seven females) NH and ten (three females) CI participants were asked to react to Positive (P), Negative (Ne), Neutral, (Nu) affective sounds virtually ending at five different distances from their body by performing fast arms flexion. Pre-motor Reaction Times (pm-RTs) were detected via EMG from postural muscles to measure action anticipation at different sound stopping distances; furthermore, the same sounds were evaluated for their level of valence and arousal perceived. Both groups showed the ability to localize the sound distances but only NH individuals modulated their pm-RTs based on the sound distance. Interestingly when the sound was not carrying affective components, as for Nu sounds, both NH and CI individuals triggered the promptest pre-motor reaction time (shorter pm-RT) when compared to P and N sounds. Only NH individuals modulated sound distance with the level of sound arousal, while sound’s valence was similarly perceived by both NH and CI individuals. These results underline the role of emotional states in action preparation and describe the specific perceptual components necessary to properly react to approaching sounds within peripersonal space.

## Introduction

We hear sounds and we deal with them continuously in our daily life; sounds carry meanings and relevant information for guiding our actions. Suppose you are interested to cross the street; you will look at the car approaching but at the same time you will pay careful attention to the sound it produces to estimate its velocity and distance for avoiding collisions. The sound produced by objects that move around us, are relevant for guiding our actions particularly in the case they are not fully visible when they are approaching us. The closer the car, the rapidly we must react and this is why the central nervous system developed a highly detailed representation for stimuli that are going to reach the nearest space around us, the so called peri-personal space (PPS) (Rizzolatti et al. 1997).

Sensory-motor information is acquired specifically in the near space: we overestimate the sound position when is reaching us (Bach et al. 2009) while the opposite happens when sounds are leaving us (Neuhoff 1998; Neuhoff 2001; Neuhoff et al. 2009; Rosenblum et al. 1993) suggesting the presence of a higher state of alert for stimuli entering the PPS. Furthermore, stimuli close to the body, both visual and auditory, enhance motor representation by modulating neural activity within the motor system (Brozzoli et al. 2010; Camponogara et al. 2015; Canzoneri et al. 2012; Finisguerra et al. 2015; Makin et al. 2009; Serino et al. 2009; Serino et al. 2011). Usually, the techniques applied to detect neural modulation within the motor system are Transcranial Magnetic Stimulation (TMS) which is now widely accepted as a tool to study the excitability of motor representation in the primary motor cortex (M1) by measuring the Motor Evoked Potentials (MEPs) amplitude (Klein-Flügge et al. 2013; Siebner et al. 2009), and Anticipatory Postural Adjustments (APAs) which are muscle activations generated by the central nervous system in a feedforward manner for anticipating actions (Massion 1992) and considered as centrally programmed commands (Aruin and Latash 1995b; Santos et al. 2010). Serino and colleagues (Serino et al. 2009) using single-pulse TMS found facilitation in the Cortico-Spinal Excitability (CSE) of hand muscles in Primary Motor Cortex (M1) while a sound was delivered close to the participant's hand but not far from it. Canzoneri and colleagues (Canzoneri et al. 2012) found that subjects were faster in detecting tactile stimuli for sounds close to hand and were faster in the presence of looming sound approaching versus sound receding. Finisguerra and colleagues (Finisguerra et al. 2015) found a larger Motor Evoked Potentials (MEPs) in right hand’s First Dorsal Interosseous (FDI) muscle by triggering optimal position in M1 for closer compared to further away approaching sounds. Camponogara and colleagues (Camponogara et al. 2015) found anticipated postural muscle activations (APAs) while reacting to approaching sounds. The special sensory-motor neural spatial map for auditory information allows auditory distance perception which is vital to detect the spatial location of environmental stimuli, and hence, regulating the action and behavior within PPS.

Natural sounds convey meanings and induce emotions that evolved to prepare individuals for efficiently contextualizing environmental events (Damasio 1998; Ekman and Davidson 1994; Lang et al. 1998; Panksepp 2004). According to the biphasic theory (Lang et al. 1997), emotions can be grouped based on their level of valence (pleasant or unpleasant) and arousal (arousing intensity). Valence, in particular, is organized around two motivational appetitive/aversive systems such as caregiving and threat (Lang et al. 1997) that developed to deal with physical survival situations (Damasio 1998; Ekman and Davidson 1994; Lang et al. 1998; Panksepp 2004). This is why emotions can be defined as “action dispositions” founded on brain states that regulate behavior along an appetitive-aversive dimension, proposing valence and arousal having a specific impact on behavior (Lang et al. 1990). In the past few years, the behavioral and neurophysiological relationships between emotion and action have been studied (Chen and Bargh 1999; Coombes et al. 2009; Hajcak et al. 2007; Rinck and Becker 2007; Schmidt et al. 2009). In behavioral studies, Chen and Bargh (1999) analyzed reaction time (RT) in individuals that asked to push or pull after the presentation on a monitor of words carrying negative or positive meanings. They found faster RTs in pushing for negative and in pulling for positive stimuli concluding that people are faster when the direction of action is congruent with the emotional content of the stimuli (Chen and Bargh 1999). In a neurophysiological study, Hajcak and colleagues (2007) found the same motor facilitation for both emotionally positive and negative stimuli compared to neutral ones (Hajcak et al. 2007), suggesting that the arousal-based facilitation in excitability of corticospinal tract is more pronounced for stimuli carrying emotional information either positive or negative compared to a neutral one. Supporting the same concept, Giovanelli et al (2013) have found higher facilitation in CSE while subjects were listening to emotionally fearful music compared to neutral and control stimulus (musical scale), while listening to music evoking happiness or sadness and displeasure presented the same CSE facilitation. Following previously mentioned researches, however, it seems there is a valence-specific facilitation in goal directed behaviors congruent to motivation along with an arousal specific pattern in neural facilitation.

Considering both the higher sensory-motor neural facilitation observed in presence of stimuli within PPS and for affective stimuli, one might expect even higher sensory-motor neural facilitation when reacting to emotional stimuli perceived within the PPS (de Haan et al. 2016; Tajadura-Jiménez et al. 2010b). Tajadura-Jiménez and colleagues studied RTs to negative and neutral images presented immediately after listening to approaching and receding sounds (Tajadura-Jiménez et al. 2010b). They found a faster reaction to the negative image compared to neutral image but more importantly approaching sounds resulted in faster RTs to both negative and neutral stimuli compared to receding sounds. Considering the valence and arousal ratings, they found that approaching sounds were more arousing than their equivalent receding sounds.

Recently it has been shown that emotional content of sounds has different impact in individuals with hearing deficits as individuals who wear cochlear implants (Ambert-Dahan et al. 2015; Paquette et al. 2018; Stabej et al. 2012; Volkova et al. 2013; Whipple et al. 2015). Cochlear implanted (CI) individuals receives sounds within a narrower dynamic range in terms of intensity and frequency, allowing lower precision in discriminating across sound intensities and frequencies (Bacon et al. 2004) but exhibit a comparable temporal resolution in detecting sound amplitude changes with Normal Hearing (NH) individuals (Kong et al. 2004; Shannon 1989; Shannon 1992). Cochlear implanted individuals, have shown the ability in perceiving the valence of sounds like discriminating happy and sad auditory stimulus, but present stronger deficit in arousal perception (Ambert-Dahan et al. 2015; Stabej et al. 2012; Volkova et al. 2013). Moreover, Ambert-Dahan and colleagues (2015) have found that CI users who received their implants after post-lingually deafness were able to discriminate different emotional valence in music (happiness, sadness, peacefulness, and threat), even in presence of deficits in arousal perception. Altogether these results indicate the important role of an intact auditory system in perceiving perceptual complexity of the sound but underlines also that cognitively individuals can compensate for hearing limitations (Ambert-Dahan et al. 2015).

In this study, we aimed at investigating the effect of dynamic affective auditory stimuli entering the PPS in modulating the motor system activity (Camponogara et al. 2015). The sounds delivered at different incremental intensities, to simulate various virtual distances within the PPS aiming to detect in details the level of promptness in reacting to sounds (Tajadura-Jiménez et al. 2010b). Furthermore, different levels of sound’s valences were considered to study the relationship between sound distance and emotional content (Chen and Bargh 1999; Coombes et al. 2009; Hajcak et al. 2007; Rinck and Becker 2007; Schmidt et al. 2009). Individuals’ modulation of APAs has been considered to quantify the facilitation in action preparation over different conditions, thus reflecting facilitation in action preparation. Furthermore, we reported the performances’ differences between CI and NH individuals, for the identification of relevant contributions through which sound perception allows efficient actions preparations (Bacon et al. 2004).

Finally, we considered and compared NH and CI individuals in three different experiments: 1) the modulation of the motor system as a function of the sound distances and emotional contents through the anticipatory adjustments 2) the behavioral accuracy in estimating the auditory distance, and 3) the individual evaluation of the sound emotional content.

## Methods

### Participants

Thirty participants were recruited. Twenty (seven females) were NH, age 23.91±3.34 years, and ten (three females) were CI, mean age 43.90±18.27 years. CI individuals presented a monoaural implant. All of the participants had no muscle-skeletal nor neurological or psychiatric history problems. The experimental protocol was approved by the members of the Ethics Committee of the Department of Neuroscience, Biomedicine, and Movement Sciences of the University of Verona. All participants gave written informed consent.

### Apparatus

All the acquisitions were performed in a sound-treated room with size 6.5×4×2.8m and the subjects positioned in its center. The background noise in the silent condition was 32dBA. The audio path was composed of an external audio-card (Scarlett 6i6, Focusrite, United Kingdom) and the resultant sounds delivered through a loudspeaker (Genelec model 8020A) mounted on a stand at 1.3m from the floor. The distance between the loudspeaker and subject was fixed at 1.5m and the seat height adjusted to align the ear of each individual with the loudspeaker woofer. Furthermore, CI individuals were sitting with the cochlear implant facing the loudspeaker while NH individuals were sitting facing the loudspeaker with the right ear facilitating attention due to the *right-ear advantage* phenomenon (Geronazzo et al. 2020). For the first experiment, APAs were detected via EMG and kinematics: two electrodes (EMG-Zero-wire system) were applied on two muscles, respectively the left and the right Erector Spinae (Aruin and Latash 1995b). To measure the kinematics, three-axial accelerometers (Analog Devices -ADXL335) were placed on the right and the left wrist. Both electrodes and accelerometers signals were recorded by an Analog to Digital Converter (CED, model Power 1401), which was remotely managed by the Spike2 software. The sampling rate was set at 5 kHz. For the second experiment the measure of the distance estimation was detected through the participant indication through upper arms movements recorded with a camera (SONY DSC-HX9V), with Full HD capabilities capturing the action through markers positioned on the body: one marker, attached to the left index finger, worked as an indicator of the position selected while two markers attached on the right arm, one on the shoulder and the other on the tip of the middle finger, were used to define the arm length (see Fig. 1). For the third experiment, a VAS score was used as explained more in detail in the next section. The use of MATLAB software allowed the high repeatability of the acquisitions.

**Fig. 1.**
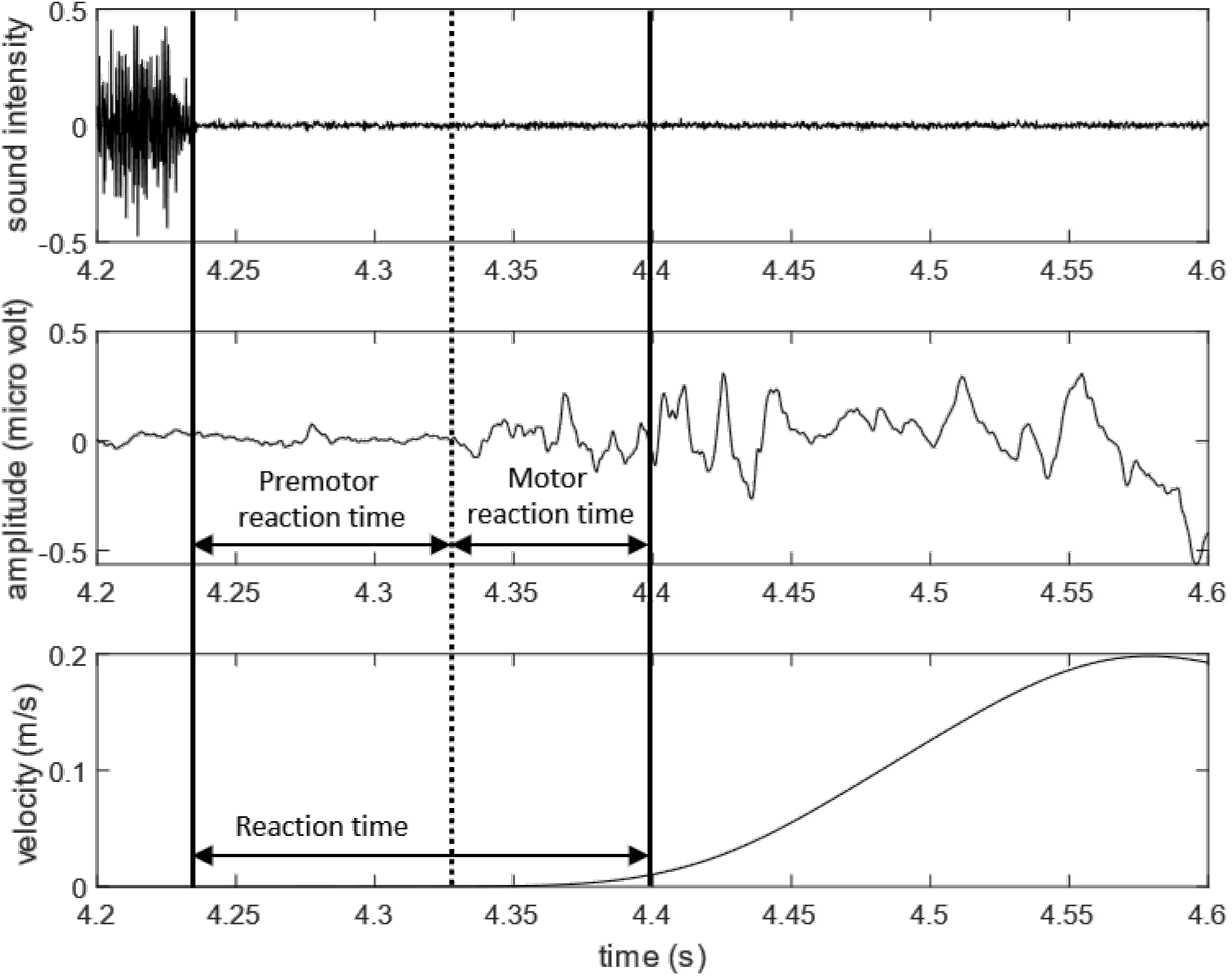
the representation of premotor reaction time, motor reaction time and reaction time in a single trial. From top to bottom: sound signal, EMG signal, and the velocity profile of the hand.

### Stimuli

Three sounds were selected. The selection was based on the International Affective Digitized Sounds (IADS) a validated library (Bradley and Lang 2007) that provides a set of standardized affective stimuli evaluated in terms of the level of valence and arousal. As presented in the list below two different samples were selected from this library (IADS identifiers are also provided) while the third stimulus was generated by MATLAB:

- id 351: applause;
- id 424: a car skidding;
- MATLAB-generated: pink noise.

The IADS samples were chosen by their values of Valence (V) and Arousal (A) scores as indicated in Table 2. The Valence was indicating the applause sound as pleasant so from now on defined as Positive (P) the car skidding as unpleasant so from now on defined as Negative (N) while the generated pink-noise is defined in the middle between the two so from now on defined neutral (Nu). To achieve uniform sound energy among stimuli, a band-pass filter with cut-off frequencies (0.25, 9.50) kHz was applied to all the samples and RMS amplitude normalization was then applied by using the Pink Noise’s level as reference. After the pre-processing, amplitude envelopes with different lengths shaped the samples to create the approaching object’s perception. The envelope was obtained by following the inverse square law between sound intensity and virtual spatial distance to the listener reference frame. The computation resulted in the so-called looming sounds (Bach et al. 2009): stimuli with exponentially rising intensity. Moreover, a 15-ms raised cosine ramp was applied to sound intensity preventing acoustic startle-reflex, and a 20ms falling cosine ramp faded out each stimulus to avoid off-responses. For each stimulus, five different ending distances were simulated with the following parameters:

- Starting sound position: 2.8 m;
- Ending sound position or nominal distance from 0.7 to 0.3 m (delta = 0.1 m; 5 dBZ);
- Approaching velocity: constant 0.7 m/s;
- Intensity range: [40, 90] dBZ.

**Table 1.** Mean and standard deviation of Valence (V) and Arousal (A) for the selected stimuli (IADS dataset).

|  |  |  | Applause | CarWreak | PinkNoise |
| --- | --- | --- | --- | --- | --- |
| IADS | V | NH | $7.32 \pm 1.62$ | $2.04 \pm 1.52$ | *NA* |
| | A | | $5.55 \pm 1.50$ | $7.99 \pm 1.66$ | *NA* |

**Table 2.**
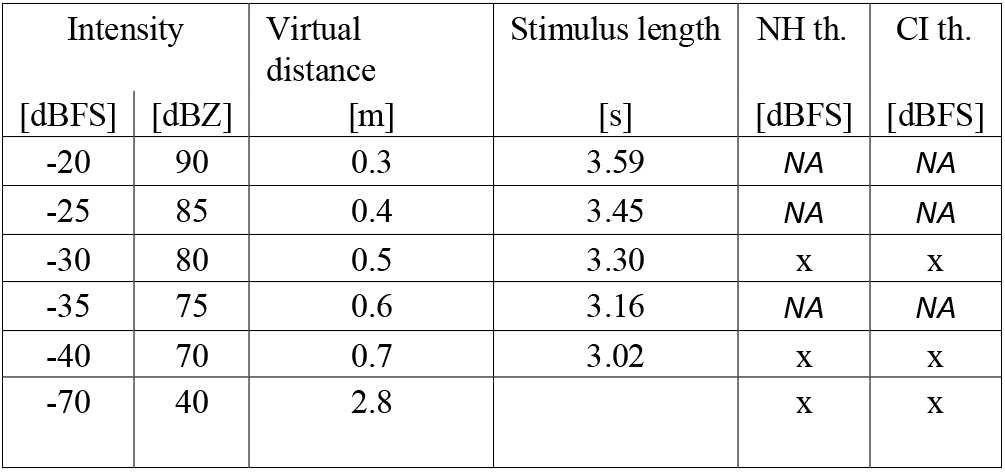
Intensity and durations of stimuli, and audiometry for PinkNoise for Normal Hearing and Cochlear Implanted.

### Procedure

#### First experiment: Anticipatory Postural Adjustment’s measurement

All participants wore an eye-mask to remove localization biases when seeing the real distance with the loudspeaker. and to avoid visual distractions and emotional effects (Tajadura-Jiménez et al. 2010a). After entering the laboratory, they were asked to sit and perform the following task: they were asked to listen to the sound by keeping a relaxed posture and, as soon as the sound stop, to raise their upper arms as fast as possible. The experiment followed a within-subjects’ design were each individual listened to 3 sounds, each of these was stopped at 5 different distances, and each distance was repeated for 5 times (5 distances × 3 sounds × 5 repetitions = 75 trials). The stimulus’s order was randomized, and a 2-minute pause was introduced after each of the three blocks made of 25 trials. A training phase allowed the subject to properly perform the task by using 12 training trials with a complex tone sound, resulting from the sum of four different complex tones of 100, 450,1.450, and 2.450 Hz (Neuhoff 1998).

The Premotor Reaction Time (pm-RT) was extracted to analyze action planning (Camponogara et al. 2015). This measure was defined as the time difference between the onset of the erector spinae muscle contraction and the ending time of the audio stimulus (Fig. 1). The onset of muscle activation was computed using the step version of the Approximated Generalized Likelihood Ratio algorithm, AGLR-step (Staude et al. 2001). The following parameters were imposed by signal inspection: sliding window of *L* =0.02*s*, the threshold for the maximum likelihood of Λ=100, the second window of *L*2 = 0.002*s* to estimate the exact onset time. The pm-RT was computed for both muscles on the left and right sides of the body. For each condition, the pm-RT values were averaged over the muscle sides and sample’s repetitions. The Reaction Time (RT) was defined as the sum of pm-RT and the Motor Reaction Time (m-RT), which accounted for the time necessary for the action execution (Camponogara et al. 2015). The m-RT was computed as the time difference between muscle activation onset and the first noticeable hand’s movement. This kinematic event was determined as the time when the velocity of the dominant finger’s reached 5% of the maximum and remained below that threshold for at least 100ms. Finally, the stimulus ending time was computed by taking the instant given by synchronous trigger.

#### Second Experiment: Distance estimation

Similar to the previous procedure, each subject was required to listen to the same previous 3 sounds organized for every 5 distances with 4 repetitions resulting in a total of 20 trials. The task required participants to indicate on their right upper arm extended forward (acting as a personal proprioceptive spatial reference for action) where the sound stopped by using the upper arm as a “meter” having the closest distance as the shoulder as a reference and the farther distance as the tip of the middle finger as a reference. To define the distance selected, as explained in the previous section, markers were used. On the right upper arm, they were located below the acromion and on the middle fingertip to define the upper arm length used as a “meter” while on the other arm the marker was located on the tip of the index finger used as a “pointer” (Fig. 2). The randomization of the stimulus’ order between groups was introduced. A pause of 2 minutes after each group allowed the subject to rest.

**Fig. 2.**
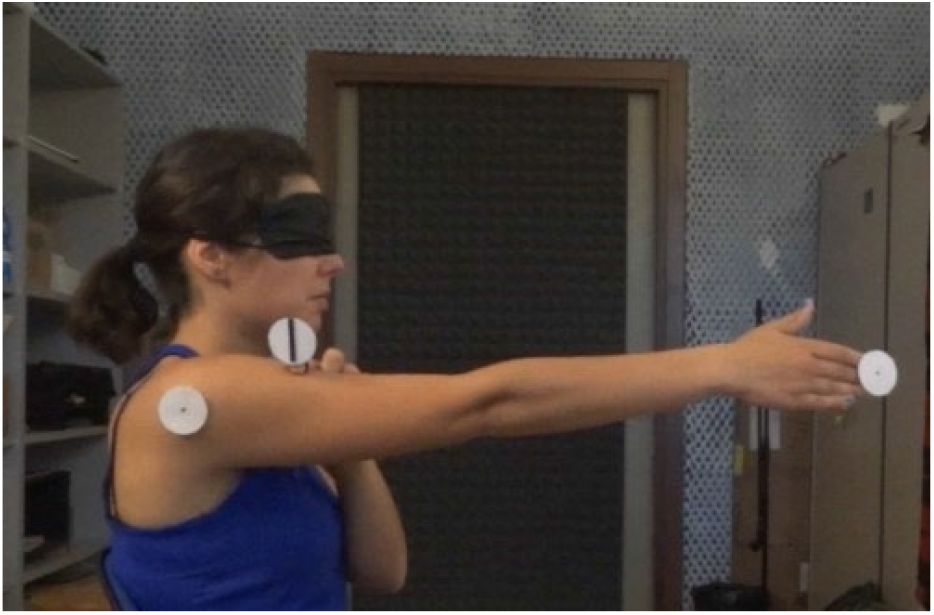
A subject while showing the perceived distance by having the right arm as meter while pointing the estimated distance with left index finger.

To compute the estimated distances a manual procedure allowed to select from the video the frame in which the subject showed the most stable position. The estimated distance was computed at that frame considering the length of the segment described by the center of the shoulder marker and the center of the pointing finger. The center of each marker was defined by a supervised algorithm and the distance between these centers calculated in pixel and then transferred to the meter scale.

#### Third Experiment: Sound Valence and Arousal evaluation

In order to assess the emotional content of the samples, the subjects were asked to rate the stimulus through the Self-Assessment Mannequin (SAM) system (Bradley and Lang 1994) delivered visually by a touchscreen laptop. For this part, the eye-mask was not used. Each participant was asked to quantify the level of valence and the level of arousal for each sound at each stopping distance. The design of stimuli delivery was the same as in the previous two experiments but only 2 repetitions were considered. Values were averaged over the repetitions.

#### Subject’s hearing sensibility evaluation

To quantify the hearing sensibility, each individual’s loudness thresholds discrimination was measured using a pink noise sound source with different sound intensity levels as reported in Table 2. The audiometric testing used the Staircase adaptive method with three forced choices (Soranzo and Grassi 2014). While normal hearing individuals showed comparable discrimination as reported in the literature (Plack and Moore 2010), the CI individuals presented large variabilities, due to the different typology and technology of their implants and due to the individual clinical situation.

### Statistics

The variables pm-RT, estimated distance, and the emotional ratings as valence and arousal have been analyzed by repeated measures ANOVA (rmANOVA) considering the Group (NH/CI) as between-subjects’ factor and Sound Valence, categorized as Positive P, Negative N, and Neutral Nu, and Distance (0.3, 0.4, 0.5, 0.6, and 0.7m) as within-subject factors. The significance level of 0.05 has been considered and the Bonferroni corrections have been applied. The SPSS software (IBM SPSS Statistics 22) has been used for statistical analysis.

## Results

### First experiment: Anticipatory Postural Adjustments

#### Premotor reaction time

The pm-RT data revealed a significant Group difference F (1,28) = 4.39, ρ = 0.045) indicating shorter pm-RTs for NH individuals (NH: 158±0.004ms, CI: 171±0.005ms). Significant main effect of Sound type has found (F (2,56) = 5.177, ρ = 0.009) showing that both groups reacted faster to Nu compared to P sound (ρ = 0.006) and slightly to N sound (ρ = 0.064); (Nu: 0.158±0.004s, N: 0.167±0.004s, P: 0.168±0.004s; Fig. 3.a and 3.b). A significant main effect of sound distance has found (F (4,112) = 9.623, ρ < 0.001) showing that both groups had longer pm-RT for the furthest distance (ρ < 0.024). Significant interaction has found for Group × Distance (F (4,112) = 4.348, ρ = 0.003) showing that NH reacted faster in 3 closest distances compared to CI (0.3m, ρ = 0.006; 0.4m, ρ = 0.046, 0.5m, ρ = 0.048). Critically, it revealed that while CI individuals did not modulate their pm-RT based on distances (Fig. 3.d), NH individuals did it (Fig. 3.c). NH individuals had shortest pm-RT in 0.3m compared to all the other distances (0.4m, ρ = 0.037; 0.5m, ρ = 0.008; 0.6m, ρ = 0.002; and 0.7, ρ <0.001), had shorter pm-RT in 0.4, 0.5, and 0.6m compared to 0.7m (ρ < 0.001). The interaction Sound × Distance didn't differentiate significantly.

**Fig. 3.**
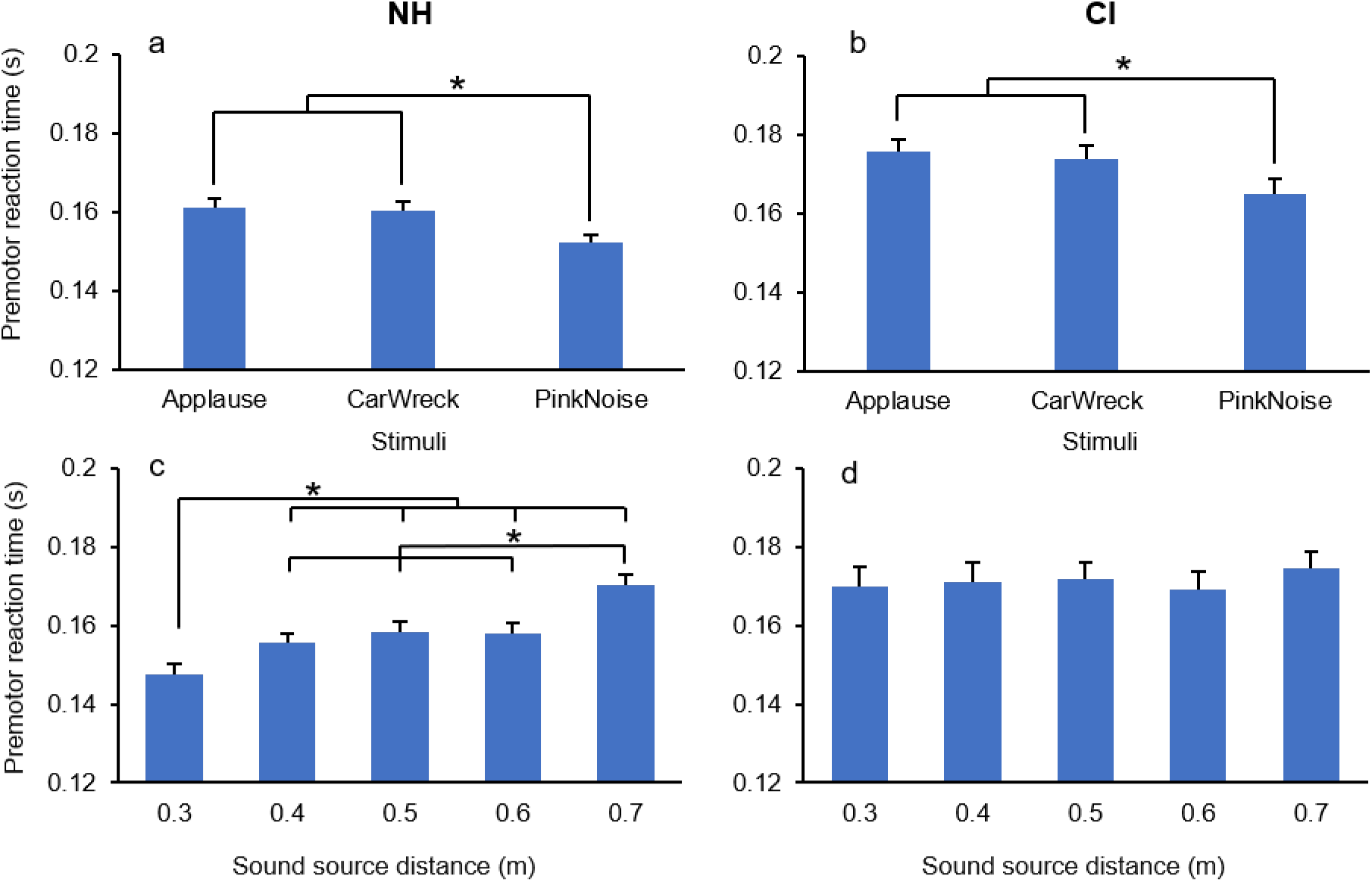
The figures on top show the pm-RT for NH (a) and CI (b) for different distances and sound types. The figures below show the pm-RT for all sound types within different distances for NH (c) and CI (d) individuals.

#### Motor reaction time

The analysis of the m-RT revealed a significant difference between groups (F (1,28) = 514.482, ρ < 0.001), showing a shorter movement time in NH individuals compared to CI individuals (NH: 0.058±0.003s, CI: 0.075±0.005s). No other significant differences have been found.

### Second Experiment: Distance estimation

rmANOVA results revealed a significant difference between groups (F (1,28) = 6.631, ρ = 0.016) showing that NH individuals perceived the sounds closer to their body compared to CI individuals (NH: 0.224±0.014m, CI: 0.286±0.020m). There was a significant main effect of distance (F (4,112) = 69.468, ρ < 0.001) and Group × Distance interaction (F (4,112) = 6.243, ρ < 0.001). NH perceived the three nearest distances (0.3, 0.4, and 0.5m) closer than CI individuals (ρ < 0.03; Fig. 4.a). Furthermore, while NH individuals discriminated all the distances (ρ < 0.001; Fig. 4.b), CI individuals could not distinguish between the two closest one (0.3m and 0.4m), but they considered three last distances as the farthest (0.5, 0.6, and 0.7m) (ρ < 0.017) while discriminating only the 0.5m as closer than the 0.7m distance (ρ = 0.027; Fig. 4c). The triple interaction Group × Sound Type × Distance was significant (F (8,224) = 2.083, ρ = 0.038). Post-hoc analysis revealed that for all the sound types, NH individuals perceived the closest distance (0.3m) significantly closer when compared with the CI (ρ = 0.032) while only for the negative sound N the same appeared for 0.4m distance (Fig. 5).

**Fig. 4.**
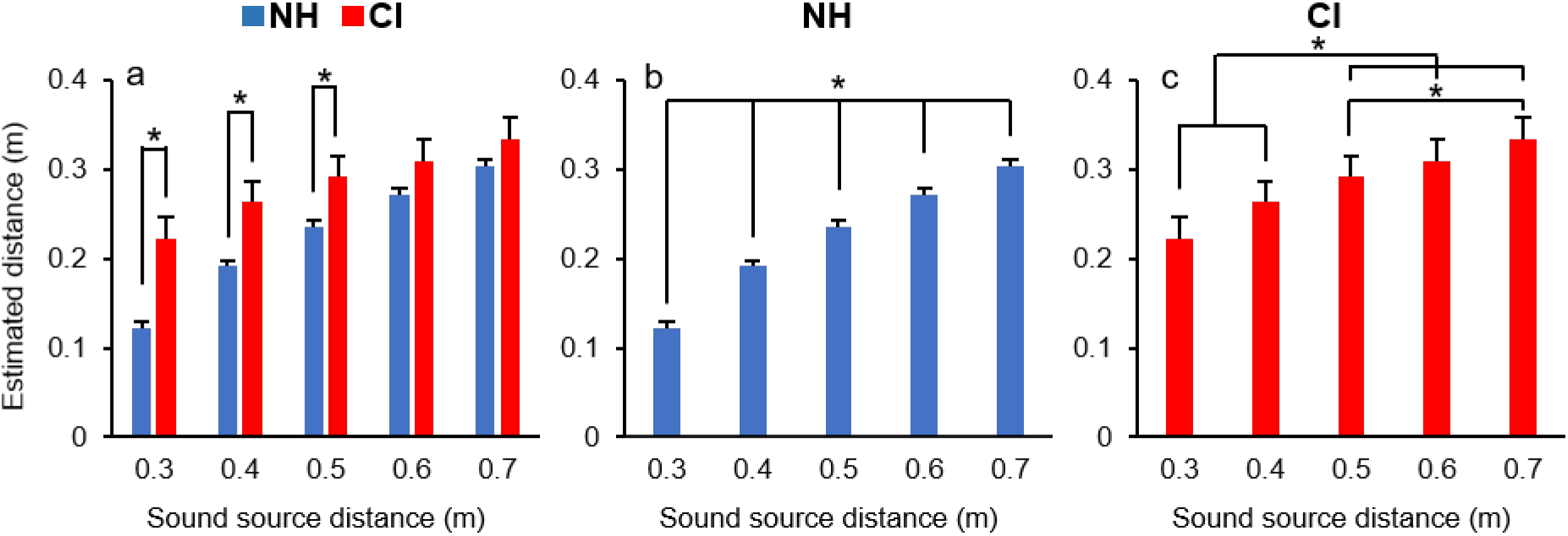
a) Estimated distances for NH (blue color) and CI (red color) groups for sound source distances. b) Estimated distances for NH c) Estimated distances for CI individuals.

**Fig. 5.**
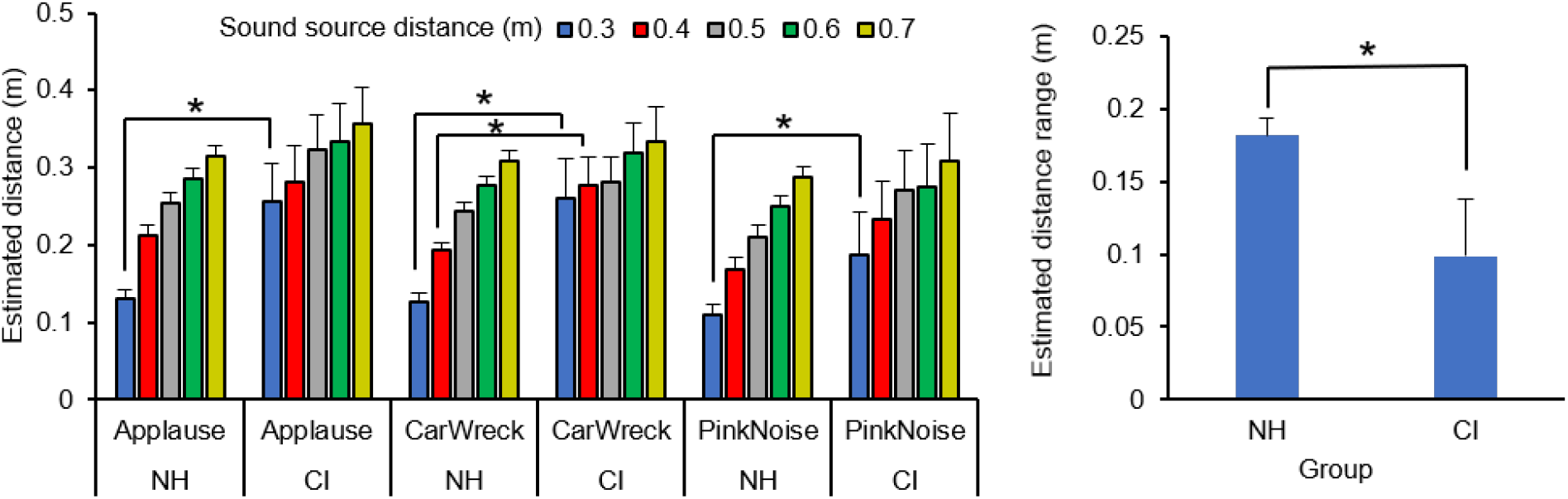
On the left estimated distance for NH and CI individuals (left) for each type of sound separately. On the right comparisons between mean estimated range distances for NH and CI individuals.

#### Range

The range utilized for discerning the distances has been analyzed by using rmANOVA and considering Group as between subjects’ factor and Sound type as a within-subject factor. The results revealed a significant difference between groups (F (1,28) = 7.165, ρ = 0.012; Fig. 5) indicating that NH individuals estimated the distances of the sounds within a wider range compared to CI individuals (NH: 0.181±0.018m, CI: 0.098±0.025m). Furthermore, results showed a significant interaction between Group × Sound type (F (2,56) = 3.299, ρ = 0.044). Post hoc analysis revealed that NH individuals located the P and N sounds within a larger range compared to CI (ρ < 0.021) while no differences occurred between the two groups for Nu sound (ρ = 0.062). Finally, CI individuals located the Nu sound within a bigger range compared to N sound (ρ = 0.014).

### Third Experiment: Sound Valence and Arousal evaluation

#### Arousal

The results showed a significant main effect of Distance (F (4,108) = 20.294, ρ < 0.001) and the interaction Group × Distance (F (4,108) = 3.303, ρ = 0.001). It should be mentioned that one CI individual was not able to complete the emotional ratings due to time limits. The post-hoc analysis showed that NH individuals perceived the closest sound (0.3m) more arousing than CI individuals (ρ = 0.027; Fig. 6.a). Critically, NH individuals modulated their arousal based on distance, while such modulation was absent in CI individuals (Figure 6).

**Fig. 6.**
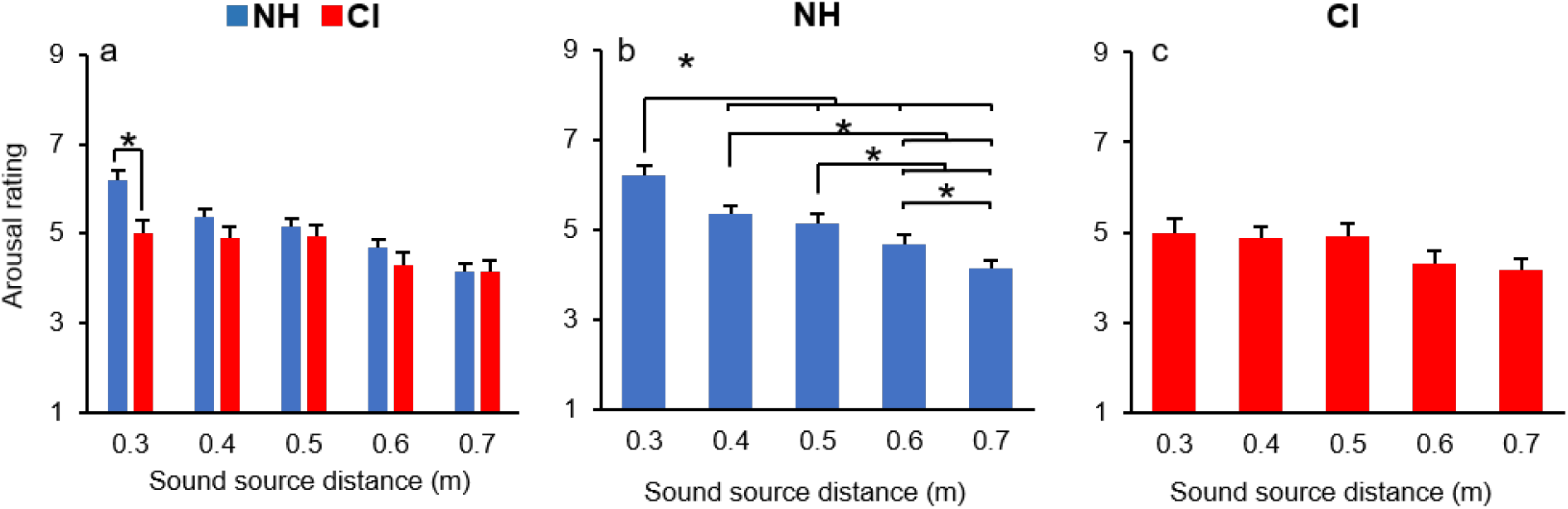
a) The arousal ratings for all the sound types for NH and CI individuals. The arousal ratings for NH (b) and CI (c) for the different distances

#### Valence

Valence was not significantly different for the two groups but the interaction Sound type × Group was significant (F (2,54) = 8.701, ρ = 0.001; Fig. 7 a). Both groups rated P higher than N (ρ = 0.001) while Nu was in between P and N (P: 5.216±0.248, N: 3.901±0.208, Nu: 4.424±0.300). Furthermore, the triple interaction Group × Sound Type × Distance was significant (F (8,216) = 2.337, ρ = 0.02) showing that NH individuals unlikely to CI individuals perceived P sound more positive than N in all different distances (ρ < 0.003; Fig. 7.b). Interestingly but only for NH group, for the closest distance 0.3m, the level of valence for Nu sound was lower than for P sound (ρ = 0.023) while for the further distances (0.5, 0.6, and 0.7m) it was higher than N sound (ρ < 0.048).

**Fig. 7.**
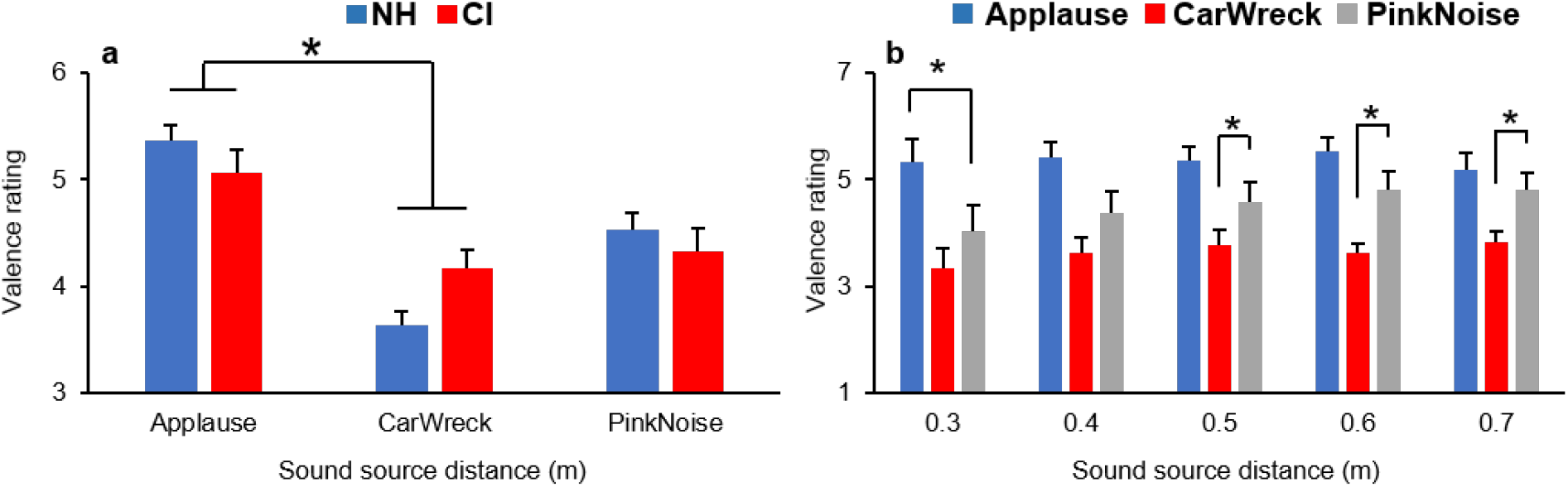
a) The valence ratings for both groups and sound types for all distances, b) the valence ratings in NH individuals for all distances and sounds

### Correlations

#### pm-RT and Estimated Distance

Linear regressions between the mean values of pm-RT and the Estimated Distance for each Sound Type and Distance have been analyzed (Figure 8). The data revealed a moderate positive correlation between pm-RT and Estimated Distance in NH individuals (*R*^2^ > 0.633) while it was negligible in CI individuals’ data mostly due to the higher variability. Crucially in NH individuals, for the P and N sounds, the correlation was stronger (P: *R*^2^ > 0.798, ρ = 0.021; N: *R*^2^ > 0.793, ρ = 0.021) and had a significantly steeper slope (P: 0.110; N: 0.120) compared to Nu sound (*R*^2^ > 0.633, ρ = 0.054; slope: 0.081) (Figure 8).

**Fig. 8.**
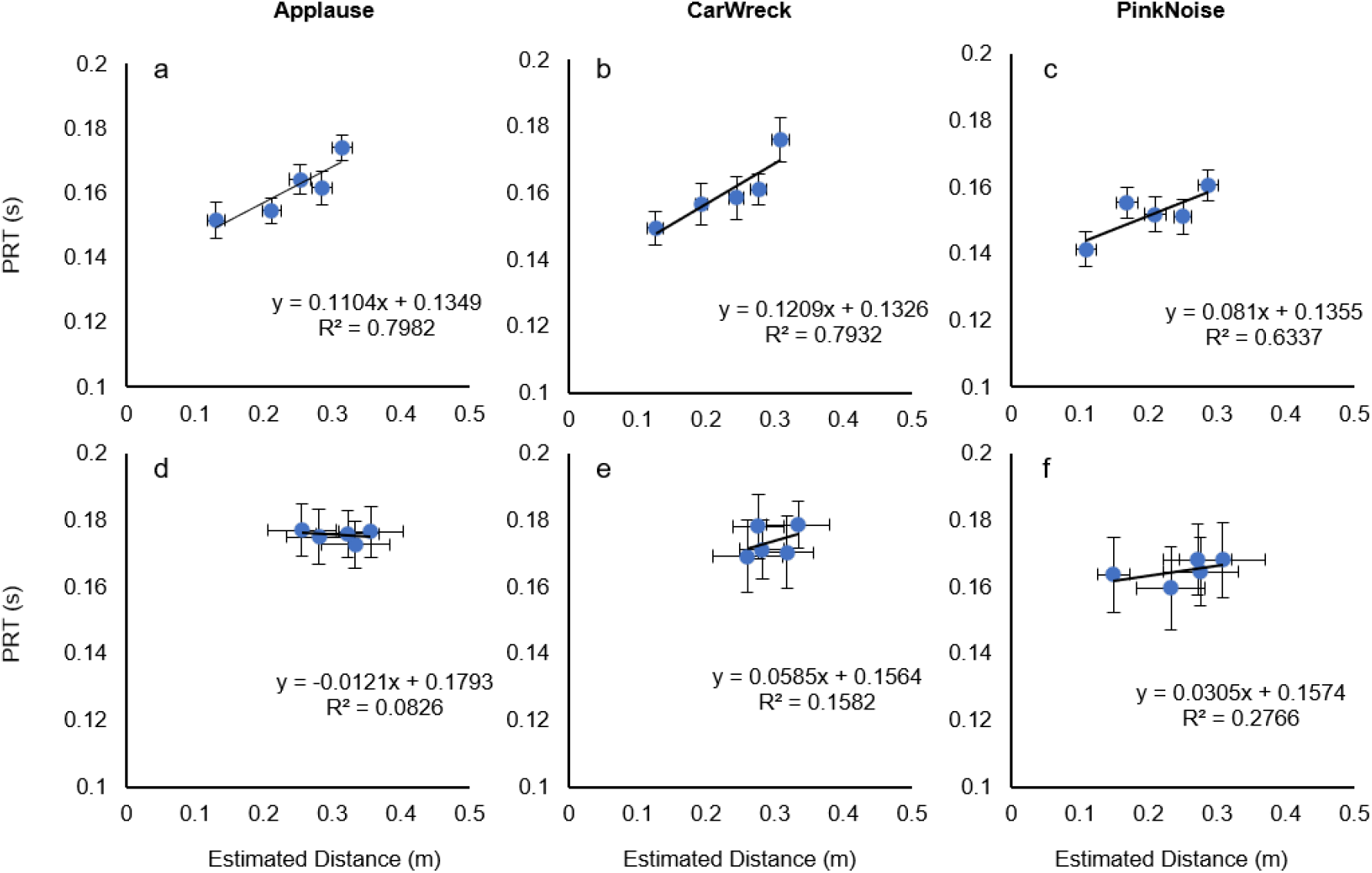
Correlation between pm-RT and Estimated Distance in NH and CI individuals for all sound types and distances. a, b, and c graphs represent NH individuals’ data and d, e, and f show the correlations in CI individuals’ data.

#### pm-RT and Arousal

The linear regressions were plotted for the mean values of pm-RT and Arousal for both NH and CI groups considering all sound types and distances (Fig. 9). The arousal data was normalized between 0 to 1. The results revealed negative correlations for all sound types in NH (*all R*^2^ > 0.667) while such a correlation was negligible in CI individuals (Fig. 9). Like the pm-RT-distance correlation in NH individuals, the correlation between pm-RT- and Arousal for the P and N sounds was stronger (P: *R*^2^ > 0.748, ρ = 0.029; N: *R*^2^ > 0.841, ρ = 0.014) and presented more inclined slopes (P: −0.082; N: −0.081) compared to Nu sound (*R*^2^ > 0.667, ρ = 0.046; slope: −0.063).

**Fig. 9.**
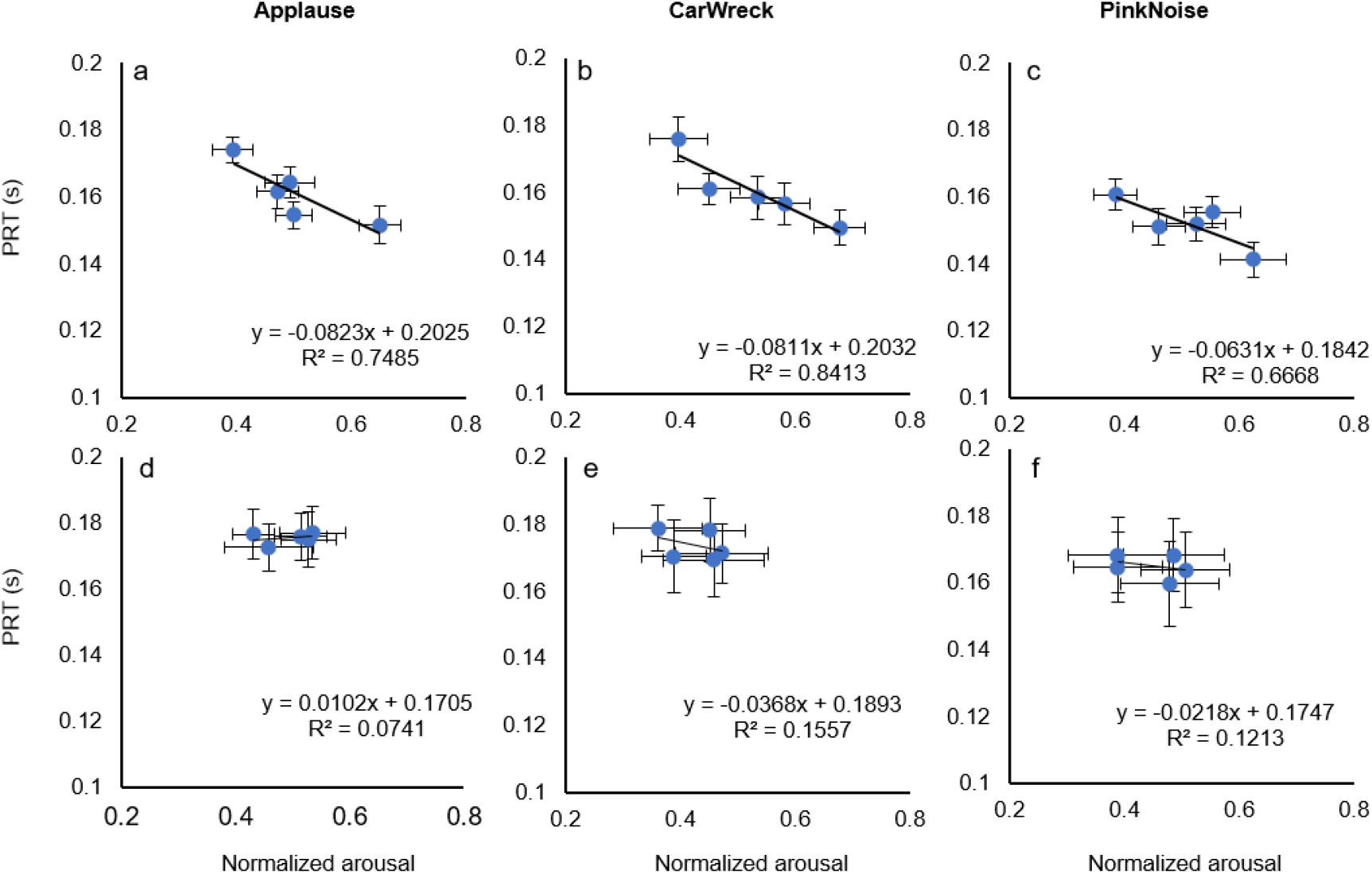
Correlation between pm-RT and normalized arousal in NH and CI individuals for all sound types and distances. a, b, and c graphs represent NH individuals’ data and d, e, and f show the correlations in CI individuals’ data.

## Discussion

In this study, we aimed at investigating the modulation of the motor system activity when dynamic affective auditory stimuli enter the individual PPS. Furthermore, to better highlight the specific dependence of the auditory information in action preparation we compared NH with CI individuals in their ability to preprogram actions. Here we showed that NH individuals were able to modulate their pm-RT as a function of the sound’s distance from the listener's ear: the closer the sound, the shorter the pm-RT. This result is in line with previous studies, indicating that when the sound source is in the near field acoustic, the motor cortex activation is enhanced (Camponogara et al. 2015; Canzoneri et al. 2012; Cléry et al. 2015; Finisguerra et al. 2015; Makin et al. 2009; Serino et al. 2009). In the same vein here, we considered pm-RT as the resultant of a feed-forward command from the motor cortex where an anticipated action is represented by an earlier activation of the motor system (Aruin and Latash 1995a; Camponogara et al. 2015; Santos et al. 2010). Our results clearly showed that normal hearing individuals finely tuned the coupling between perception and action for anticipating their reactions of about 10 ms for every 10 cm of sound stopping closer to their body. CI individuals, on the contrary, presented a constant pm-RTs across all the sounds’ distances indicating their inability to modulate properly their timing for adequate action preparation. This lack of modulation in action preparation suggests the necessity to elaborate alternative strategies for maintaining an acceptable level of safety while navigating in the surrounding space (Fletcher et al. 2019a; Fletcher et al. 2019b; Huang et al. 2019; Huang et al. 2017). Interestingly when data of pm-RT were analyzed based on the type of sound, both NH and CI individuals were reacting faster for neutral compared to positive and negative sounds. A possible interpretation is that while positive and negative sounds convey meanings that require time for being decoded and recognized, on the contrary, in neutral (Nu) sounds meaning is absent allowing for a prompt pre-motor reaction. Indeed, our results showed that both groups NH and CI participants prepared the action (pm-RT) by using the same strategy: shorter pm-RT for neutral (Nu) compared to positive (P) and negative (N) sounds (Beatty et al. 2016). Interestingly when the time of movement was considered (from the first muscle activation to the first upper arm displacement captured by the kinematics) no modulation of muscle activation was present, nor for normal hearing and cochlear implanted individuals. This result is particularly relevant for NH in indicating that for an efficient reaction to sounds, decisions to move should be taken way before action initiation.

As far as for the ability to localize sounds in space, NH indicated the stop of sounds closer than CI did. This happened for the three closest sounds (the 0.3m 0.4m and 0.5m). Moreover, NH compared to CI individuals located the sounds within a significantly larger spatial range for positive and negative sounds (the range was defined by the closer and the farther points of localization). This might be caused by the reduced abilities of CI hearing system in discriminating sound intensities and frequencies both highly relevant for distance estimations (Brungart 1999; Courtois et al. 2019; Kopčo and Shinn-Cunningham 2011; Zahorik 2002). Interestingly though for the neutral sound both NH and CI selected the same range. The Neutral sound contains indeed the same amount of information in the frequency bandwidth of interest so that it is easier to be processed from both NH and CI individuals.

Nevertheless, CI individuals still showed a certain ability to localize sounds’ distances even in presence of impairments in the auditory cues (Bacon et al. 2004; Busby and Clark 2000; Zeng and Galvin III 1999): how then they were able to localize where the sounds stopped? One possible interpretation is that CI individuals relied mostly on the temporal component of the approaching sounds. Indeed, CI individuals have reported having normal or better than normal temporal resolution in amplitude changes detection measured by gap valuation and amplitude modulation detection tasks (Kong et al. 2004; Shannon 1989; Shannon 1992).

As far as for evaluating the emotional content of the sounds delivered, for NH individuals the closer the sound was, the more arousing was evaluated; it is worth notice that such modulation was not present in CI individuals. The result obtained for NH individuals is in line with the literature since it has been shown that approaching sounds are perceived as a warning signal and as a consequence localized closer to the body as they normally are (Neuhoff 1998; Neuhoff 2001; Neuhoff et al. 2009; Rosenblum et al. 1993; Schiff and Oldak 1990), in a similar vein, these stimuli have been found to facilitate the action (Bach et al. 2009; Finisguerra et al. 2015; Marinovic et al. 2014; Serino et al. 2009). This perceptual-motor modulations signaling higher state of alert for approaching sounds reflects the presence of a defensive mechanism, showing the tendency of keeping the body far from the potential threats and keeping threats possibly outside the PPS (Neuhoff 1998; Neuhoff 2016; Vagnoni et al. 2017; Vagnoni et al. 2012). It is worth to note that the emotional content of sounds affects distance evaluation as well: the closer the sound is, the higher the perceived arousal (Tajadura-Jiménez et al. 2009). Here we showed that while this is the case for NH, it is not for CI individuals (Ambert-Dahan et al. 2015; Paquette et al. 2018). Critically, the level of arousal has found to be responsible in emotion-driven changes in the cortico-spinal tract (CST) excitability and neural activity within the motor system (Baumgartner et al. 2007; Coombes et al. 2009; Hajcak et al. 2007), here we found that CI individuals being not able to activate properly their level of arousal they do lack the proper modulation in action preparation (pm-RT). As shown in the literature, the low spectral resolution provided by the cochlear implants is a relevant deficit that compromises the assessment of the emotional content in sounds (Busby and Clark 2000; Handel 1995; Paquette et al. 2018).

Both NH and CI distinguished the sounds’ valences: The P sound has been evaluated more positive than N sound and the Nu sound was evaluated in between the P and N sounds. Although a detailed perception of differences among sounds in CI individuals is impaired (Ambert-Dahan et al. 2015; Paquette et al. 2018; Stabej et al. 2012; Volkova et al. 2013; Whipple et al. 2015), they were still able to discriminate happy and sad emotional auditory stimulus above chance. Even though the level of precision of CI in such discrimination is not comparable with NH individuals (Stabej et al. 2012; Volkova et al. 2013) Ambert-Dahan and colleagues (2015) have found that for CI who received their implants after post-lingual deafness presented a remarkable ability in discriminating different emotional valences in music.

NH individuals presented a strong positive correlation between pm-RT and sound distance, demonstrating their fine-tuning in the perception-action relationship (Camponogara et al. 2015). This is in line with previous studies, indicating modifications in action preparation along with the level of brain activation for stimuli within the PPS (Camponogara et al. 2015; Canzoneri et al. 2012; Fogassi et al. 1996; Tajadura-Jiménez et al. 2010b). Also, NH presented a strong negative correlation between pm-RT and Arousal; a higher level of arousal led to a prompter reaction. Already in previous studies, the level of arousal was positively correlated with the increment in the sensory-motor neural correlates (Coombes et al. 2009; Hajcak et al. 2007; Schmidt et al. 2009), here we add that the ability to localize sound in the surrounding space is positively correlated with the level of arousal and the promptness in action preparation.

Based on the results, it can be suggested that action preparation for reacting to stimuli within the auditory PPS, relies on the emotional content of the stimuli along with their distance. Interestingly, the data represented a stronger correlation with a steeper ramp for emotional sounds compared to Nu sound in NH individuals, suggesting a more fine-tuning when there is a higher state of alert (Bach et al. 2009; Beatty et al. 2016; Vagnoni et al. 2012). It is known that sensory-motor facilitations within the PPS are originated by defensive and goal-directed purposes evoked by threatening and or pleasant stimuli (Coombes et al. 2009; De Vignemont and Iannetti 2015; Graziano and Cooke 2006; Lane and Nadel 2002; Lang et al. 1998).

The effect of multisensory neural representations in PPS considering approaching affective stimuli might emphasize on amygdala pivotal role (Anderson and Sobel 2003; Sabatinelli et al. 2005; Small et al. 2003). Approaching sounds have found to be perceived as warning signals, recruiting a distributed neural network including the motor and premotor cortices, the intraparietal sulcus, and the amygdala (Bach et al. 2007). Furthermore, the amygdala seems to play also a pivotal role in the definition of the PPS. In a seminal study, Kennedy and colleagues (Kennedy et al. 2009) reported the case of a patient with a complete amygdala lesion who was lacking any sense of personal space. This result has been corroborated by an imaging study showing activation of the amygdala related to close personal proximity (Kennedy et al. 2009). Critically, it is shown that the action modulations within PPS strongly correlates with the sense of personal space, as it significantly changes when the PPS perception changes; for example due to tool use (Bassolino et al. 2010; Canzoneri et al. 2013; Maravita and Iriki 2004; Serino et al. 2007) or different phobias (de Haan et al. 2016; Taffou and Viaud-Delmon 2014). In this regard, it can be concluded that the absence of modulation in action anticipation in CI individuals for sounds stopping at different distances within the PPS, might be partially due to the lack of arousal perception.

In conclusion, having an intact hearing system is necessary for an accurate perceptual elaboration of the characteristics of the sound as intensity and spectral frequency distribution, leading to properly interact within the environment full of predictable and non-predictable sound stimuli. In the same vein, perceiving precisely the content of auditory stimuli is also crucial to correctly modulate preparatory actions as a function of sound distance; this ability is limited in CI individuals being constrained by their artificial hearing that provides capabilities of the specific type of cochlear implant (Ambert-Dahan et al. 2015; Paquette et al. 2018; Stabej et al. 2012; Volkova et al. 2013; Whipple et al. 2015). To our knowledge, this is the first study that measures the action-perception capabilities in CI individuals. By considering the departure in the results obtained combining NH with CI individuals, in this study, we were able to better underline the importance of having an intact hearing system of sound perception for action preparation. Finally, it is important to mention that our results suggest that improving the technology of cochlear implants, will improve not just speech recognition (Nie et al. 2006; van Hoesel and Tyler 2003) or music perception (Drennan and Rubinstein 2008; Gfeller et al. 2006; McDermott 2004), but also action preparation abilities in individuals with cochlear implants addressing also the everyday interaction with the external environment.

## References

Ambert-Dahan E, Giraud A-L, Sterkers O, Samson S (2015) Judgment of musical emotions after cochlear implantation in adults with progressive deafness Frontiers in psychology 6:181

Anderson AK, Sobel N (2003) Dissociating intensity from valence as sensory inputs to emotion Neuron 39:581–583

Aruin AS, Latash ML (1995a) Directional specificity of postural muscles in feed-forward postural reactions during fast voluntary arm movements Experimental Brain Research 103:323–332

Aruin AS, Latash ML (1995b) The role of motor action in anticipatory postural adjustments studied with self-induced and externally triggered perturbations Experimental Brain Research 106:291–300

Bach DR, Neuhoff JG, Perrig W, Seifritz E (2009) Looming sounds as warning signals: The function of motion cues International Journal of Psychophysiology 74:28–33

Bach DR et al. (2007) Rising sound intensity: an intrinsic warning cue activating the amygdala Cerebral Cortex 18:145–150

Bacon SP, Fay RR, Popper AN (2004) Compression: from cochlea to cochlear implants. Springer,

Bassolino M, Serino A, Ubaldi S, Làdavas E (2010) Everyday use of the computer mouse extends peripersonal space representation Neuropsychologia 48:803–811

Baumgartner T, Willi M, Jäncke L (2007) Modulation of corticospinal activity by strong emotions evoked by pictures and classical music: a transcranial magnetic stimulation study Neuroreport 18:261–265

Beatty GF, Cranley NM, Carnaby G, Janelle CM (2016) Emotions predictably modify response times in the initiation of human motor actions: A meta-analytic review Emotion 16:237

Bradley MM, Lang PJ (1994) Measuring emotion: the self-assessment manikin and the semantic differential Journal of behavior therapy and experimental psychiatry 25:49–59

Bradley MM, Lang PJ (2007) The International Affective Digitized Sounds (; IADS-2): Affective ratings of sounds and instruction manual University of Florida, Gainesville, FL, Tech Rep B-3

Brozzoli C, Cardinali L, Pavani F, Farnè A (2010) Action-specific remapping of peripersonal space Neuropsychologia 48:796–802

Brungart DS (1999) Auditory localization of nearby sources. III. Stimulus effects The Journal of the Acoustical Society of America 106:3589–3602

Busby P, Clark GM (2000) Pitch estimation by early-deafened subjects using a multiple-electrode cochlear implant The Journal of the Acoustical Society of America 107:547–558

Camponogara I, Komeilipoor N, Cesari P (2015) When distance matters: Perceptual bias and behavioral response for approaching sounds in peripersonal and extrapersonal space Neuroscience 304:101–108

Canzoneri E, Magosso E, Serino A (2012) Dynamic sounds capture the boundaries of peripersonal space representation in humans PloS one 7:e44306

Canzoneri E, Ubaldi S, Rastelli V, Finisguerra A, Bassolino M, Serino A (2013) Tool-use reshapes the boundaries of body and peripersonal space representations Experimental Brain Research 228:25–42

Chen M, Bargh JA (1999) Consequences of automatic evaluation: Immediate behavioral predispositions to approach or avoid the stimulus Personality and social psychology bulletin 25:215–224

Cléry J, Guipponi O, Wardak C, Hamed SB (2015) Neuronal bases of peripersonal and extrapersonal spaces, their plasticity and their dynamics: knowns and unknowns Neuropsychologia 70:313–326

Coombes SA, Tandonnet C, Fujiyama H, Janelle CM, Cauraugh JH, Summers JJ (2009) Emotion and motor preparation: a transcranial magnetic stimulation study of corticospinal motor tract excitability Cognitive, Affective, & Behavioral Neuroscience 9:380–388

Courtois G, Grimaldi V, Lissek H, Estoppey P, Georganti E (2019) Perception of Auditory Distance in Normal-Hearing and Moderate-to-Profound Hearing-Impaired Listeners Trends in Hearing 23:2331216519887615 doi:10.1177/2331216519887615

Damasio AR (1998) Emotion in the perspective of an integrated nervous system Brain research reviews 26:83–86

de Haan AM, Smit M, Van der Stigchel S, Dijkerman HC (2016) Approaching threat modulates visuotactile interactions in peripersonal space Experimental Brain Research 234:1875–1884

De Vignemont F, Iannetti G (2015) How many peripersonal spaces? Neuropsychologia 70:327–334

Drennan WR, Rubinstein JT (2008) Music perception in cochlear implant users and its relationship with psychophysical capabilities Journal of rehabilitation research and development 45:779

Ekman PE, Davidson RJ (1994) The nature of emotion: Fundamental questions. Oxford University Press,

Finisguerra A, Canzoneri E, Serino A, Pozzo T, Bassolino M (2015) Moving sounds within the peripersonal space modulate the motor system Neuropsychologia 70:421–428

Fletcher MD, Hadeedi A, Goehring T, Mills SR (2019a) Electro-haptic enhancement of speech-in-noise performance in cochlear implant users Scientific reports 9:1–8

Fletcher MD, Hadeedi A, Goehring T, Mills SR (2019b) Electro-haptic hearing: Speech-in-noise performance in cochlear implant users is enhanced by tactile stimulation of the wrists

Fogassi L, Gallese V, Fadiga L, Luppino G, Matelli M, Rizzolatti G (1996) Coding of peripersonal space in inferior premotor cortex (area F4) Journal of neurophysiology 76:141–157

Geronazzo M, Barumerli R, Cesari P (2020) Motor planning unveils the shape of human auditory peripersonal space: an immersive virtual reality approach IEEE Transactions on Human-Machine Systems (under review)

Gfeller KE, Olszewski C, Turner C, Gantz B, Oleson J (2006) Music perception with cochlear implants and residual hearing Audiology and Neurotology 11:12–15

Graziano MS, Cooke DF (2006) Parieto-frontal interactions, personal space, and defensive behavior Neuropsychologia 44:845–859

Hajcak G, Molnar C, George MS, Bolger K, Koola J, Nahas Z (2007) Emotion facilitates action: a transcranial magnetic stimulation study of motor cortex excitability during picture viewing Psychophysiology 44:91–97

Handel S (1995) Chapter 12 - Timbre Perception and Auditory Object Identification. In: Moore BCJ (ed) Hearing. Academic Press, San Diego, pp 425–461. doi:https://doi.org/10.1016/B978-012505626-7/50014-5

Huang J, Lu T, Sheffield B, Zeng F-G (2019) Electro-Tactile Stimulation Enhances Cochlear-Implant Melody Recognition: Effects of Rhythm and Musical Training Ear and hearing Publish Ahead of Print doi:10.1097/aud.0000000000000749

Huang J, Sheffield B, Lin P, Zeng F-G (2017) Electro-tactile stimulation enhances cochlear implant speech recognition in noise Scientific reports 7:2196

Kennedy DP, Gläscher J, Tyszka JM, Adolphs R (2009) Personal space regulation by the human amygdala Nature neuroscience 12:1226

Klein-Flügge MC, Nobbs D, Pitcher JB, Bestmann S (2013) Variability of human corticospinal excitability tracks the state of action preparation Journal of Neuroscience 33:5564–5572

Kong Y-Y, Cruz R, Jones JA, Zeng F-G (2004) Music perception with temporal cues in acoustic and electric hearing Ear and hearing 25:173–185

Kopčo N, Shinn-Cunningham BG (2011) Effect of stimulus spectrum on distance perception for nearby sources The Journal of the Acoustical Society of America 130:1530–1541

Lane RD, Nadel L (2002) Cognitive neuroscience of emotion. Oxford University Press,

Lang PJ, Bradley MM, Cuthbert BN (1990) Emotion, attention, and the startle reflex Psychological review 97:377

Lang PJ, Bradley MM, Cuthbert BN (1997) Motivated attention: Affect, activation, and action Attention and orienting: Sensory and motivational processes 97:135

Lang PJ, Bradley MM, Cuthbert BN (1998) Emotion, motivation, and anxiety: Brain mechanisms and psychophysiology Biological psychiatry 44:1248–1263

Makin TR, Holmes NP, Brozzoli C, Rossetti Y, Farne A (2009) Coding of visual space during motor preparation: approaching objects rapidly modulate corticospinal excitability in hand-centered coordinates Journal of Neuroscience 29:11841–11851

Maravita A, Iriki A (2004) Tools for the body (schema) Trends in cognitive sciences 8:79–86

Marinovic W, Tresilian JR, de Rugy A, Sidhu S, Riek S (2014) Corticospinal modulation induced by sounds depends on action preparedness The Journal of physiology 592:153–169

Massion J (1992) Movement, posture and equilibrium: interaction and coordination Progress in neurobiology 38:35–56

McDermott HJ (2004) Music perception with cochlear implants: a review Trends in amplification 8:49–82

Neuhoff JG (1998) Perceptual bias for rising tones Nature 395:123

Neuhoff JG (2001) An adaptive bias in the perception of looming auditory motion Ecological Psychology 13:87–110

Neuhoff JG (2016) Looming sounds are perceived as faster than receding sounds Cognitive research: principles and implications 1:15

Neuhoff JG, Planisek R, Seifritz E (2009) Adaptive sex differences in auditory motion perception: Looming sounds are special Journal of Experimental Psychology: Human Perception and Performance 35:225

Nie K, Barco A, Zeng F-G (2006) Spectral and temporal cues in cochlear implant speech perception Ear and hearing 27:208–217

Panksepp J (2004) Affective neuroscience: The foundations of human and animal emotions. Oxford university press,

Paquette S, Ahmed G, Goffi-Gomez M, Hoshino A, Peretz I, Lehmann A (2018) Musical and vocal emotion perception for cochlear implants users Hearing research 370:272–282

Plack CJ, Moore DR (2010) The oxford handbook of auditory science: hearing vol 3. Oxford University Press New York, NY,

Rinck M, Becker ES (2007) Approach and avoidance in fear of spiders Journal of behavior therapy and experimental psychiatry 38:105–120

Rizzolatti G, Fadiga L, Fogassi L, Gallese V (1997) The space around us Science 277:190–191

Rosenblum LD, Wuestefeld AP, Saldana HM (1993) Auditory looming perception: Influences on anticipatory judgments Perception 22:1467–1482

Sabatinelli D, Bradley MM, Fitzsimmons JR, Lang PJ (2005) Parallel amygdala and inferotemporal activation reflect emotional intensity and fear relevance Neuroimage 24:1265–1270

Santos MJ, Kanekar N, Aruin AS (2010) The role of anticipatory postural adjustments in compensatory control of posture: 2. Biomechanical analysis Journal of Electromyography and Kinesiology 20:398–405

Schiff W, Oldak R (1990) Accuracy of judging time to arrival: effects of modality, trajectory, and gender Journal of Experimental Psychology: Human Perception and Performance 16:303

Schmidt L, Cléry-Melin M-L, Lafargue G, Valabrègue R, Fossati P, Dubois B, Pessiglione M (2009) Get aroused and be stronger: emotional facilitation of physical effort in the human brain Journal of Neuroscience 29:9450–9457

Serino A, Annella L, Avenanti A (2009) Motor properties of peripersonal space in humans PloS one 4:e6582

Serino A, Bassolino M, Farne A, Ladavas E (2007) Extended multisensory space in blind cane users Psychological science 18:642–648

Serino A, Canzoneri E, Avenanti A (2011) Fronto-parietal areas necessary for a multisensory representation of peripersonal space in humans: an rTMS study Journal of cognitive neuroscience 23:2956–2967

Shannon RV (1989) Detection of gaps in sinusoids and pulse trains by patients with cochlear implants The Journal of the Acoustical Society of America 85:2587–2592

Shannon RV (1992) Temporal modulation transfer functions in patients with cochlear implants The Journal of the Acoustical Society of America 91:2156–2164

Siebner HR, Hartwigsen G, Kassuba T, Rothwell JC (2009) How does transcranial magnetic stimulation modify neuronal activity in the brain? Implications for studies of cognition Cortex 45:1035–1042

Small DM, Gregory MD, Mak YE, Gitelman D, Mesulam MM, Parrish T (2003) Dissociation of neural representation of intensity and affective valuation in human gustation Neuron 39:701–711

Soranzo A, Grassi M (2014) PSYCHOACOUSTICS: a comprehensive MATLAB toolbox for auditory testing Frontiers in psychology 5:712

Stabej KK, Smid L, Gros A, Zargi M, Kosir A, Vatovec J (2012) The music perception abilities of prelingually deaf children with cochlear implants International journal of pediatric otorhinolaryngology 76:1392–1400

Staude G, Flachenecker C, Daumer M, Wolf W (2001) Onset detection in surface electromyographic signals: a systematic comparison of methods EURASIP Journal on Advances in Signal Processing 2001:867853

Taffou M, Viaud-Delmon I (2014) Cynophobic fear adaptively extends peri-personal space Frontiers in psychiatry 5:122

Tajadura-Jiménez A, Kitagawa N, Väljamäe A, Zampini M, Murray MM, Spence C (2009) Auditory–somatosensory multisensory interactions are spatially modulated by stimulated body surface and acoustic spectra Neuropsychologia 47:195–203

Tajadura-Jiménez A, Larsson P, Väljamäe A, Västfjäll D, Kleiner M (2010a) When room size matters: acoustic influences on emotional responses to sounds Emotion 10:416

Tajadura-Jiménez A, Väljamäe A, Asutay E, Västfjäll D (2010b) Embodied auditory perception: The emotional impact of approaching and receding sound sources Emotion 10:216

Vagnoni E, Andreanidou V, Lourenco SF, Longo MR (2017) Action ability modulates time-to-collision judgments Experimental Brain Research 235:2729–2739

Vagnoni E, Lourenco SF, Longo MR (2012) Threat modulates perception of looming visual stimuli Current biology 22:R826–R827

van Hoesel RJ, Tyler RS (2003) Speech perception, localization, and lateralization with bilateral cochlear implants The Journal of the Acoustical Society of America 113:1617–1630

Volkova A, Trehub SE, Schellenberg EG, Papsin BC, Gordon KA (2013) Children with bilateral cochlear implants identify emotion in speech and music Cochlear Implants International 14:80–91

Whipple CM, Gfeller K, Driscoll V, Oleson J, McGregor K (2015) Do communication disorders extend to musical messages? An answer from children with hearing loss or autism spectrum disorders Journal of music therapy 52:78–116

Zahorik P (2002) Assessing auditory distance perception using virtual acoustics The Journal of the Acoustical Society of America 111:1832–1846

Zeng F-G, Galvin III JJ (1999) Amplitude mapping and phoneme recognition in cochlear implant listeners Ear and hearing 20:60–74

